# IL-13 and IL-17A Activate β1 Integrin through an NF-kB/Rho kinase/PIP5K1γ pathway to Enhance Force Transmission in Airway Smooth Muscle

**DOI:** 10.1101/2024.05.01.592042

**Authors:** Uyen Ngo, Ying Shi, Prescott Woodruff, Kevan Shokat, William DeGrado, Hyunil Jo, Dean Sheppard, Aparna B. Sundaram

**Affiliations:** Division of Pulmonary, Critical Care, Allergy and Sleep, Department of Medicine, University of California, San Francisco, California, USA; Sandler Asthma Basic Research Center, University of California, San Francisco, California, USA; Department of Cellular and Molecular Pharmacology, University of California, San Francisco, California, USA; Howard Hughes Medical Institute, University of California, San Francisco, California, USA; Cardiovascular Research Institute, University of California, San Francisco, California, USA; Department of Pharmaceutical Chemistry, University of California, San Francisco, California, USA

## Abstract

Integrin activation resulting in enhanced adhesion to the extracellular matrix plays a key role in fundamental cellular processes. Although G-protein coupled receptor-mediated integrin activation has been extensively studied in non-adherent migratory cells such as leukocytes and platelets, much less is known about the regulation and functional impact of integrin activation in adherent stationary cells such as airway smooth muscle. Here we show that two different asthmagenic cytokines, IL-13 and IL-17A, activate type I and IL-17 cytokine receptor families respectively, to enhance adhesion of muscle to the matrix. These cytokines also induce activation of β1 integrins as detected by the conformation-specific antibody HUTS-4. Moreover, HUTS-4 binding is significantly increased in the smooth muscle of patients with asthma compared to healthy controls, suggesting a disease-relevant role for aberrant integrin activation. Indeed, we find integrin activation induced by a β1 activating antibody, the divalent cation manganese, or the synthetic peptide β1-CHAMP, dramatically enhances force transmission in collagen gels, mouse tracheal rings, and human bronchial rings even in the absence of cytokines. We further demonstrate that cytokine-induced activation of β1 integrins is regulated by a common pathway of NF-kB-mediated induction of RhoA and its effector Rho kinase, which in turn stimulates PIP5K1γ-mediated synthesis of PIP_2_ resulting in β1 integrin activation. Taken together, these data identify a previously unknown pathway by which type I and IL-17 cytokine receptor family stimulation induces functionally relevant β1 integrin activation in adherent smooth muscle and help explain the exaggerated force transmission that characterizes chronic airways diseases such as asthma.

**SIGNIFICANCE STATEMENT:** Integrin activation plays a central role in regulating cellular adhesion and migration. While chemokine-mediated integrin activation has been extensively studied in circulating cells, the role and impact of other cytokine families on non-migratory cells remains incompletely characterized. Here, we demonstrate in airway smooth muscle that asthmagenic cytokines IL-13 and IL-17A stimulate type I and IL-17 cytokine receptor families to induce β1 integrin activation and enhance adhesion. We also identify a common pathway linking NF-kB/RhoA/Rho kinase with PIP5K1γ/PIP2/β1 integrin activation. We show that airway biopsies from asthmatic patients have increased active β1 integrin staining in the muscle, and furthermore that β1 integrin activation alone dramatically enhances force transmission, underscoring the disease-relevant impact of cytokine-mediated integrin activation in adherent muscle.

## INTRODUCTION

Allergic asthma is a chronic and heterogeneous disorder affecting the airways. Airway smooth muscle plays an integral role in the pathobiology of asthma by regulating airway tone as well as contributing to airway inflammation and remodeling (1). These processes are stimulated by inflammatory mediators such as cytokines, chemokines, and growth factors. The pro-inflammatory Th2 cytokine IL-13 plays a central role in the pathogenicity of asthma by augmenting airway smooth muscle hyperresponsiveness and remodeling (2, 3). Th17 cytokines such as IL-17A have also been shown to influence the severity of airway hyperresponsiveness (4, 5).

Integrins are heterodimeric transmembrane proteins that play a central role in adhesion, as well as regulation of migration, survival, and growth. One of the hallmarks of integrin physiology is the ability to engage in bidirectional signaling, where cues from the intracellular or extracellular environment can trigger cellular responses (6). Our prior work showed that inhibiting the association of integrin α5β1 with its ligand fibronectin or integrin α2β1 with collagen I could inhibit the exaggerated force transmission induced by IL-13 and IL-17A in contracting airway smooth muscle without altering actin-myosin cross-bridge cycling (7, 8). Curiously, integrin ligation has no effect on baseline force transmission in the absence of pro-inflammatory cytokines. Given the importance of cytokines in driving the hyperresponsiveness that is a central cause of morbidity and mortality in chronic airway diseases such as asthma, a better understanding of how cytokines modulate integrin-dependent adhesion of airway smooth muscle is needed.

The adhesive function of integrins is modulated by their activation state. This has been best studied in circulating cells where integrins are normally maintained in a partially (9) or fully bent closed conformation (10) to keep cells from inappropriately adhering. In response to chemokines and other activators of G protein-coupled receptors (such as thrombin or ATP, in the case of platelets), integrins undergo a rapid and dramatic conformational change, assuming the extended and open conformation (11) required for binding to integrin ligands and firm adhesion. However, in adherent cells where it is assumed integrins are already bound to the underlying matrix, the extent of integrin activation and functional relevance of activation state remain important unanswered questions.

In this study, we demonstrate that the asthmagenic cytokines IL-13 and IL-17A activate type I and IL-17 cytokine receptor families in airway smooth muscle resulting in an unexpected activation of surface β1 integrins. We identify that IL-13 and IL-17A induce nuclear translocation of NF-kB resulting in increased expression of RhoA and its downstream effector, Rho kinase. Rho kinase in turn stimulates PIP5K1γ-mediated synthesis of PIP_2_ to recruit talin resulting in activation of β1 integrins in airway smooth muscle. Furthermore, we show differential activation of β1 integrins in the airway smooth muscle of asthmatics compared to healthy controls. We couple this with disease-relevant implications by establishing that integrin activation alone is sufficient to enhance force transmission in contracting smooth muscle even in the absence of pro-inflammatory cytokines.

## RESULTS

### IL-13 and IL-17A activate integrins to enhance adhesion of airway smooth muscle to extracellular ligands

Integrins play an important role in tethering cells to the surrounding extracellular environment. We previously found that ligation of integrins α2β1 and α5β1 with function-blocking antibodies completely prevented adhesion of human airway smooth muscle (HASM) cells to collagen I and fibronectin, respectively (7, 8). Blockade of either integrin also abrogated the enhanced force transmission induced by IL-13 and IL-17A ex vivo and respiratory resistance in allergic asthma models in vivo by impairing tethering of muscle to the surrounding extracellular matrix. Notably, there were no effects of integrin inhibition on baseline responses in the absence of cytokines.

Integrins transition between low- and high-affinity ligand binding states, where the integrin adopts a bent-closed or extended-open conformation, respectively. While integrin activation in circulating cells is tightly controlled, integrin activation in adherent cells has been less well characterized in part because the extended-open conformation is thought to predominate. Incubation of primary airway smooth muscle cells with IL-13 did not affect surface expression of either integrin α5β1 or α2β1 (7, 8), but did increase expression of an epitope on the integrin β1 subunit accessible only when the integrin is in the extended-open confirmation (12). Furthermore, treatment with IL-13 increased adhesion of smooth muscle to the α2β1 ligand, collagen I, suggesting IL-13 increases force transmission by activating the α2β1 integrin on airway smooth muscle cells.

To determine whether integrin activation is functionally important for adhesion to collagen I and fibronectin, and whether integrin activation could be a generalized response to multiple asthmagenic cytokines, we performed adhesion assays with HASM cells treated with either IL-13 or IL-17A. We found cells treated with the canonical Th2 cytokine IL-13 had increased adhesion to both collagen I and fibronectin compared to untreated controls (**Fig. 1a**), with the degree of adhesion depending on concentration of ligand applied to the surface. The pro-inflammatory cytokine IL-17A is associated with the neutrophilic inflammatory response and development of severe forms of asthma. Treatment with IL-17A also increased adhesion of HASM cells to a range of surface concentrations of collagen I and fibronectin compared to untreated controls (**Fig. 1b**). Corresponding to these changes in adhesive strength, treatment of HASM cells with IL-13 or IL-17A increased expression of the active integrin β1 subunit compared to vehicle as detected by the conformation-specific antibody HUTS-4, which recognizes the extended-open conformation of the β1 subunit (12). The divalent cation Mn^2+^, which alters the conformation of the extracellular domain of integrins to induce activation (13), was used as a positive control (**Fig. 1c, Supplemental Fig. 1a-b**). These findings support the conclusion that IL-13 and IL-17A both activate α5β1 and α2β1 integrins.

**Figure 1:**
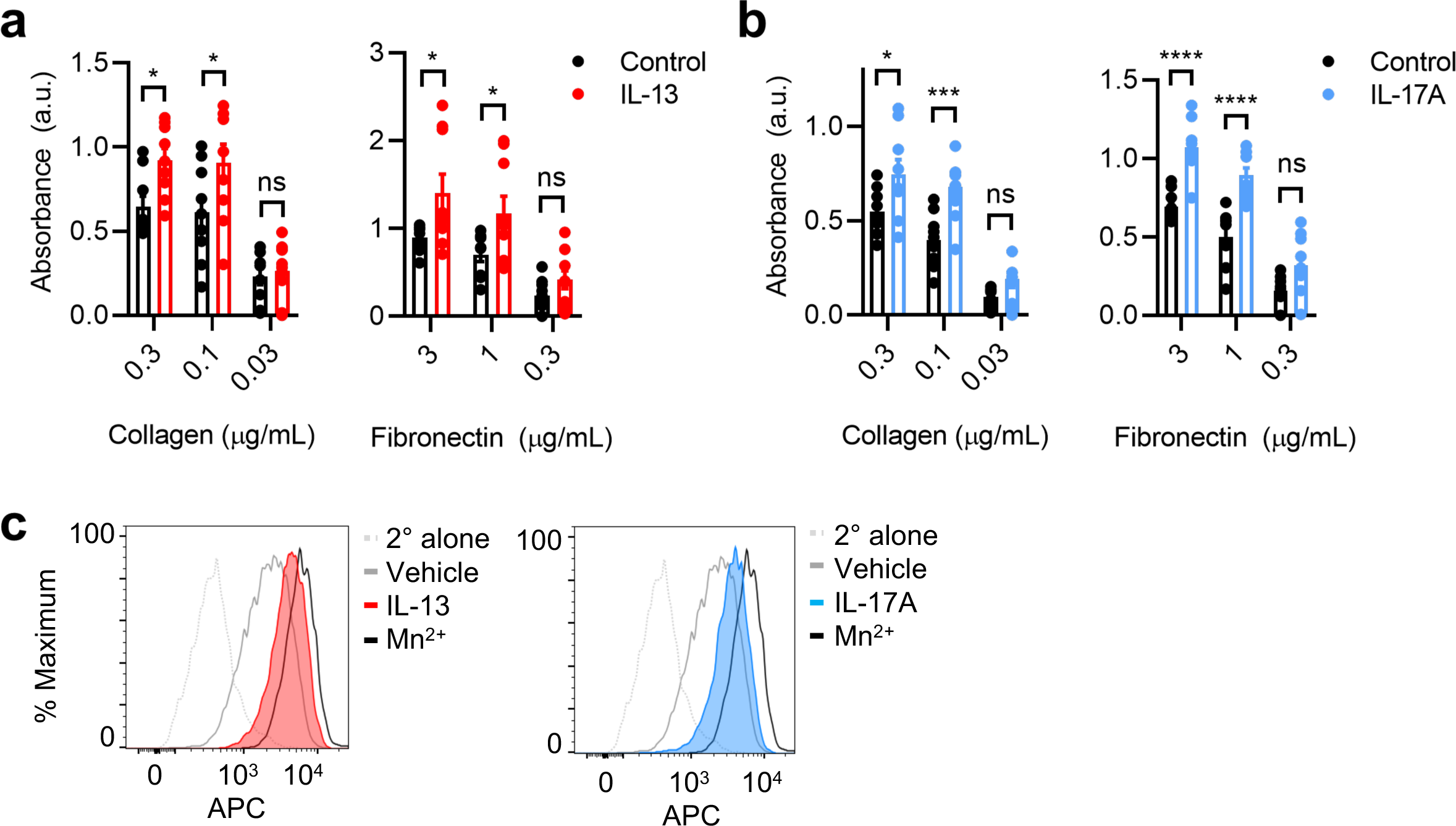
IL-13 and IL-17A enhance adhesion and activate β1 integrins. **a-b,** Adhesion as measured by absorbance of crystal violet at 595 nm of HASM cells to various concentrations of collagen and fibronectin after treatment with vehicle, IL-13 (100 ng/mL), or IL-17A (100 ng/mL) for 12 hours. Experiments performed in triplicate with 3 biological replicates for all panels. **c,** Representative histograms for active β1 integrin in HASM cells treated with vehicle, IL-13 (100 ng/mL), IL-17A (100 ng/mL) for 12 hours, or Mn^2+^ (1 mM) for 20 minutes followed by labeling with antibody specific for active β1 integrin (HUTS-4) and secondary (2°) conjugated to allophycocyanin (APC). Results from 3 biological replicates shown in Supplemental Figure 1. Data are mean ± s.e.m. for **a-b**. 2-way ANOVA with Sidak’s multiple-comparison test. *P<0.05, ***P<0.001, ****P<0.0001, ns=not significant.

### Airway smooth muscle integrins are activated in human asthma

To determine whether these in vitro observations had relevance to behavior of smooth muscle integrins in human tissue, bronchial rings taken from donor lungs were treated with vehicle, IL-13, or Mn^2+^ followed by staining for the active β1 integrin with HUTS-4. Compared to vehicle, exposure to IL-13 increased active β1 integrin staining in airway smooth muscle to a degree comparable with Mn^2+^ (**Fig. 2a**). To further understand the relevance of these findings in chronic airways disease, we obtained de-identified airway biopsy samples from three patients with asthma and three healthy non-smoking controls (**Supplemental Table 1**). We found increased active β1 integrin staining in the airway smooth muscle of asthma patients compared to healthy non-smoking controls (**Fig. 2b**). This supports the conclusion that cytokine-induced activation of integrins occurs in airway smooth muscle in situ and has disease-relevance in asthma.

**Figure 2:**
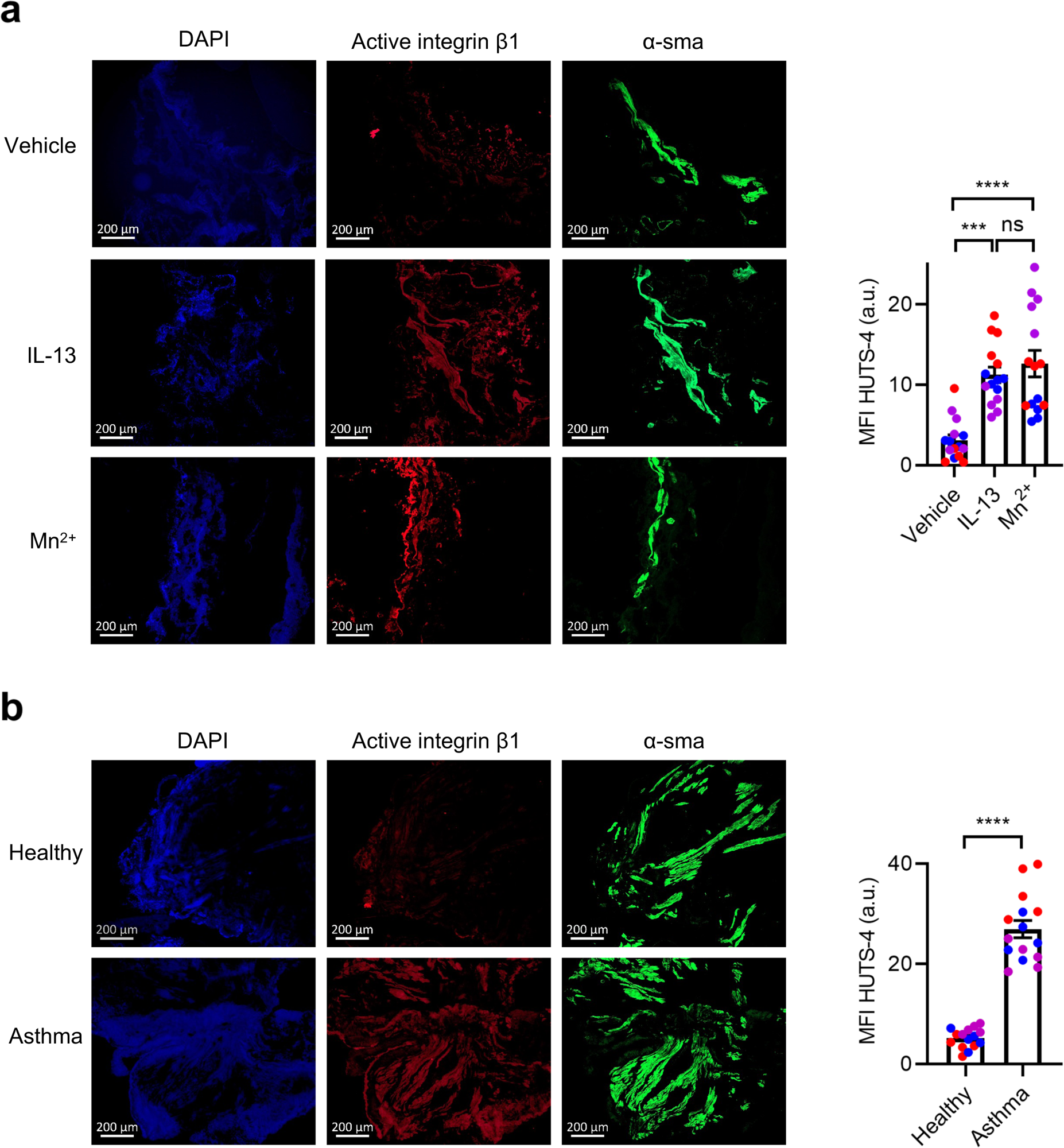
Smooth muscle integrins are activated by cytokines in asthma. **a,** Representative immunofluorescence from human bronchial rings treated with vehicle, IL-13 (100 ng/mL) for 12 hours, or Mn^2+^ (1 mM) for 2 hours, then sectioned and stained with antibodies specific for active β1 integrin (HUTS-4, red), anti-α-smooth muscle actin (α-sma, green), and DAPI (blue). Scale bar = 200 μm. Median fluorescence intensity (MFI) of HUTS-4 was measured in five sections per sample (n=3 donors per group). **b,** Representative immunofluorescence from endobronchial airway biopsy specimens from asthma and healthy non-smokers stained with antibodies specific for active β1 integrin (HUTS-4, red), anti-α-smooth muscle actin (green), and DAPI (blue). Scale bar = 200 μm. MFI of HUTS-4 was measured in five sections per sample (n=3 donors per group). Data are mean ± s.e.m. for **a-b**. 2-way ANOVA with Tukey’s multiple-comparison test for **a**. 2-tailed t-test for **b**. ***P<0.001, ****P<0.0001, ns=not significant.

### Integrin activation is sufficient to enhance force transmission by airway smooth muscle

Integrins can be activated exogenously by the addition of activating antibodies, metal ions, or synthetic peptides. To determine whether integrin activation is sufficient to enhance force transmission in airway smooth muscle, even in the absence of asthmagenic cytokines, we evaluated the contractile response to histamine in HASM cells embedded in a collagen gel matrix incubated with the β1 integrin activating antibody TS2/16 or IgG control for 12 hours. Integrin activation resulted in enhanced contraction of collagen gels compared to IgG control. Furthermore, when HASM cells treated with TS2/16 were exposed to c15, a small molecule inhibitor of integrin α2β1, the exaggerated contraction was eliminated (**Fig. 3a**). HASM cells treated with Mn^2+^ produced similar results (**Fig. 3b**).

**Figure 3:**
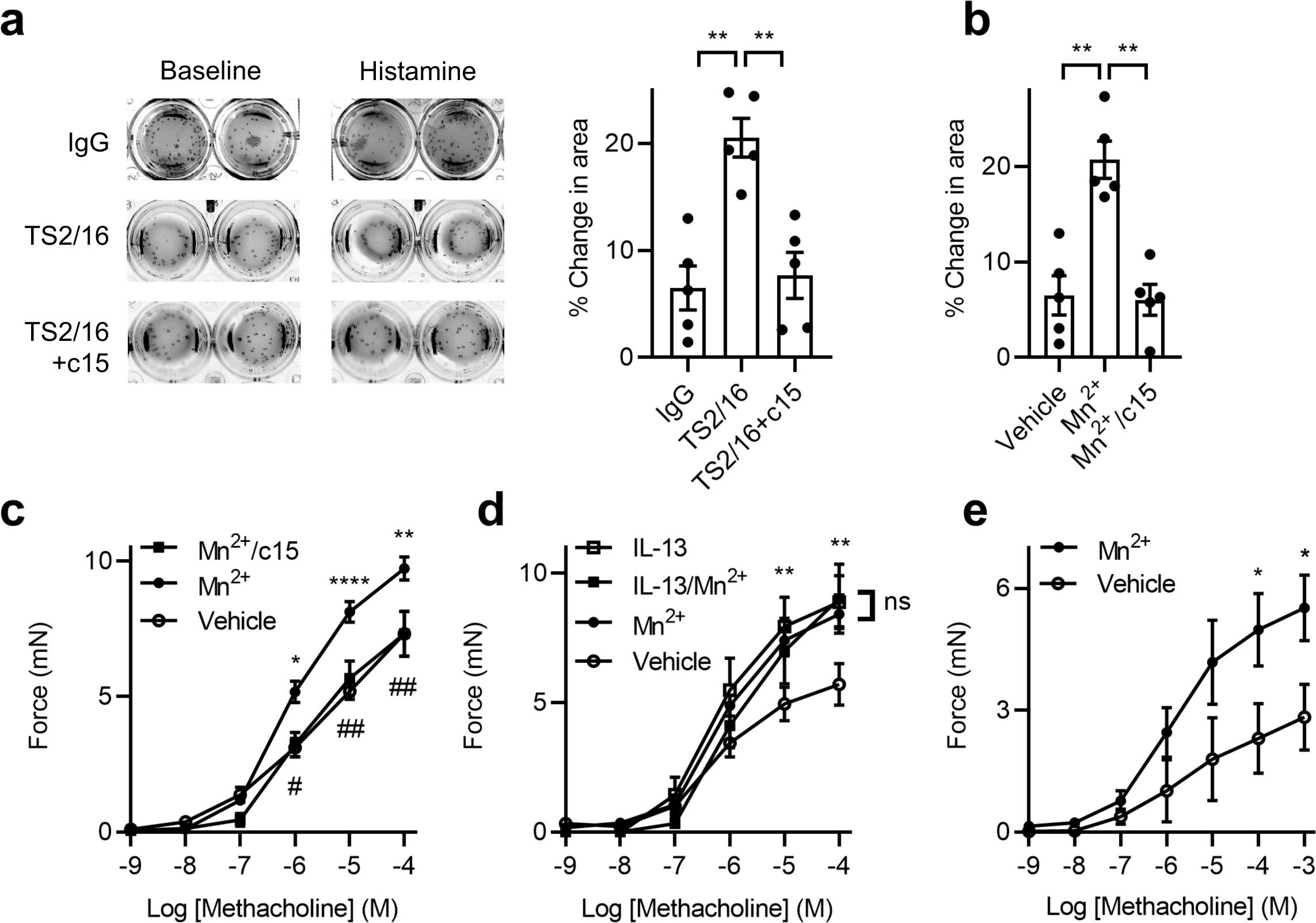
Integrin activation enhances force transmission. **a-b,** Representative images and quantification of percentage change in area after histamine 100 μM relative to baseline after HASM embedded in collagen gels are exposed to (**a**) IgG control, β1 activating antibody (TS2/16, 10 μM) for 12 hours, or (**b**) vehicle, Mn^2+^ (1 mM) for 20 minutes, followed by the α2 integrin inhibitor c15 (20 μM) for 1 hour. n=5 samples per group. **c,** Contractile force measured in mouse tracheal rings after incubation with vehicle or Mn^2+^ (1 mM) for 2 hours, followed by the α2 integrin inhibitor c15 (20 μM) for 1 hour, with a range of concentrations of methacholine. n=5-7 rings per group. **d,** Contractile force measured in mouse tracheal rings after incubation with vehicle or IL-13 (100 ng/mL) for 12 hours, followed by Mn^2+^ (1 mM) for 2 hours, with a range of concentrations of methacholine. n=3-9 rings per group. **e,** Contractile force measured in human bronchial rings after incubation with vehicle or Mn^2+^ (1 mM) for 2 hours, with a range of concentrations of methacholine. n=6 rings per group. Data are mean ± s.e.m. for **a-e**. 2-way ANOVA with Tukey’s multiple-comparison test for **a-b**. 2-way ANOVA with repeated measures, Tukey’s multiple-comparison test for **c-e**. *P<0.05, **P<0.01, ****P<0.0001, ns=not significant. #P<0.05, ##P<0.01 vs Mn^2+^ for **c**.

Given the paucity of β1 integrin activating antibodies targeting mice and the difficultly of adequate antibody penetration into tissue, we assessed contractile responses to methacholine in mouse tracheal rings incubated with Mn^2+^ or vehicle for 2 hours. External activation of integrins with Mn^2+^ enhanced force transmission compared to vehicle. Notably, incubation of Mn^2+^ treated rings with c15 eliminated the exaggerated force transmission, confirming that Mn^2+^ effects on force transmission are integrin-dependent (**Fig. 3c**). Because the effects of Mn^2+^ are local and do not induce separation of integrin transmembrane domains required for bidirectional signaling (14, 15), we confirmed our findings using the synthetic peptide β1-CHAMP that targets the transmembrane helix of the β1 subunit to activate integrins in a manner mimicking the physiologic activator talin (**Supplemental Fig. 2a**) (16, 17). Treatment with β1-CHAMP also enhanced force transmission in mouse tracheal rings compared to vehicle (**Supplemental Fig. 2b**). Interestingly, while mouse tracheal rings exposed to the pro-inflammatory cytokine IL-13 had exaggerated force transmission as expected, the addition of Mn^2+^ to IL-13 treated rings did not further augment force transmission compared to treatment with either IL-13 or Mn^2+^ alone (**Fig. 3d**). The ability of Mn^2+^ to recapitulate, but not augment, the effect of IL-13 supports the hypothesis that cytokines induce inside-out signaling changes resulting in integrin activation and increased adhesion to extracellular matrix ligands. Human bronchial rings treated with Mn^2+^ also had similar effects to mouse tracheal rings, with an increase in force transmission in response to methacholine compared to vehicle (**Fig. 3e**). Taken together, these findings support the conclusion that integrin activation is sufficient to enhance force transmission in airway smooth muscle and provide a compelling explanation for the exaggerated force transmission induced by asthmagenic cytokines.

### Integrin activation by asthmagenic cytokines depends on Rho kinase-induced activation of PIP5K1γ and PIP_2_

Physiologic activation of integrins is a coordinated process that involves multiple proteins that converge on the cytoskeletal protein talin which serves as a linker that couples the actin cytoskeleton to the cytoplasmic tail of the β-integrin subunit. Talin attachment is spatially directed by the plasma membrane lipid phosphatidylinositol 4,5-bisphosphate (PIP_2_) (18, 19), which disrupts the auto-inhibitory form of talin to expose its integrin binding site (20, 21). To confirm that this pathway is involved in β1 integrin activation in airway smooth muscle, we treated HASM cells with a cell-permeable form of PIP_2_ (diC16-PIP_2_) or its carrier vehicle alone and found diC16-PIP_2_ increased active β1 integrin by flow cytometry (**Fig. 4a**). Similarly, diC16-PIP_2_ enhanced force transmission in mouse tracheal rings in response to increasing doses of methacholine compared to rings treated with the carrier vehicle alone (**Fig. 4b**).

**Figure 4:**
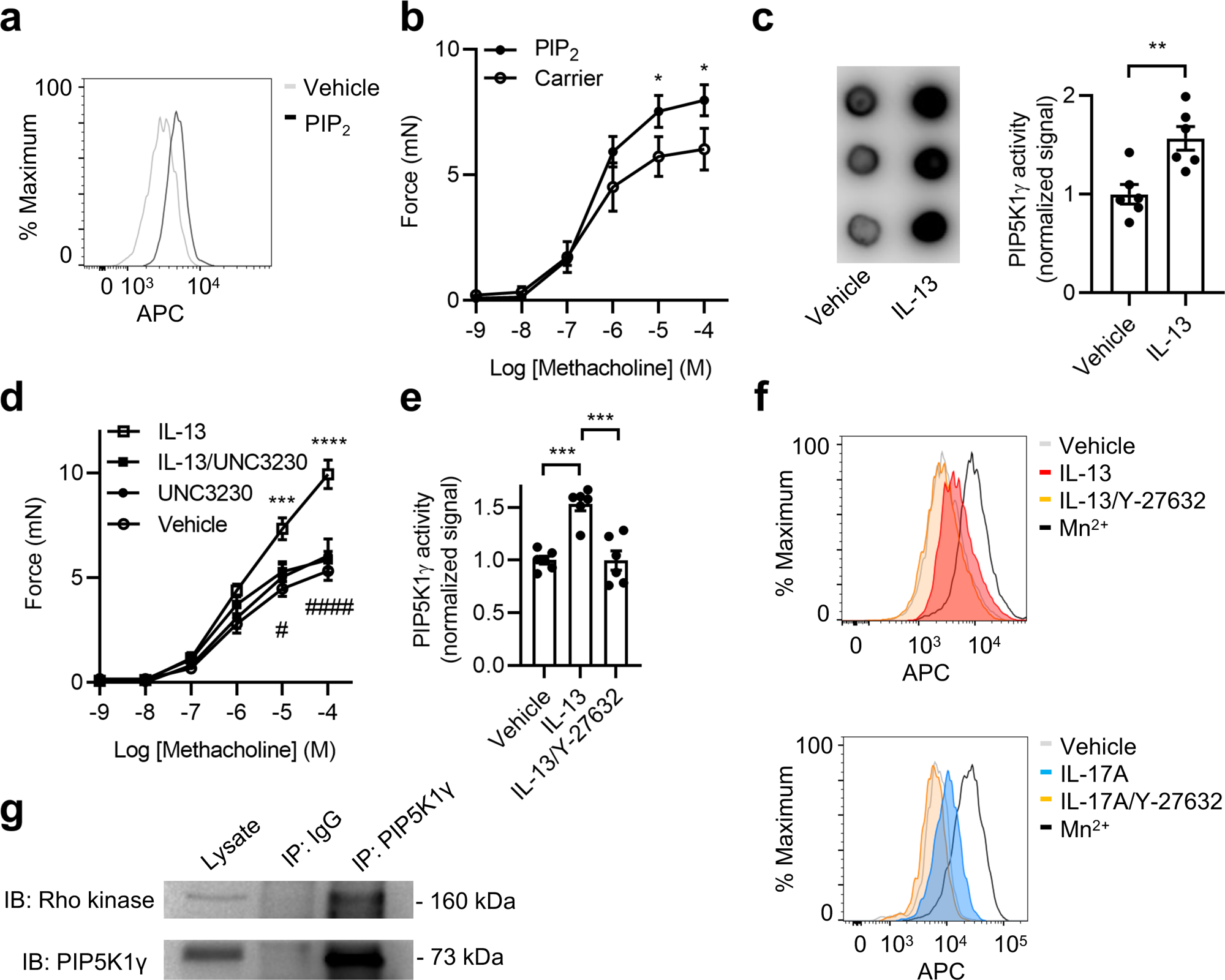
Integrin activation depends on Rho kinase-dependent activation of PIP5K1γ and PIP_2_. **a,** Representative histogram for active β1 integrin in HASM cells treated with lipid carrier 1 in the presence or absence of diC16-PIP_2_ (10 μM) for 1 hour, followed by labeling with antibody specific for active β1 integrin (HUTS-4) and secondary conjugated to allophycocyanin (APC). **b,** Contractile force measured in mouse tracheal rings after incubation with lipid carrier 1 in the presence or absence of diC16-PIP_2_ (10 μM) for 1 hour with a range of concentrations of methacholine. n=6-10 rings per group. **c,** Representative blot and quantification of PIP5K1γ activity normalized to vehicle as measured by transfer of [γ-^32^P] to PI4P from [γ-^32^P] ATP in HASM after treatment with vehicle or IL-13 (100 ng/mL) for 12 hours followed by lysis and IP with anti-PIP5K1γ antibody. n=6 biological replicates per group. **d,** Contractile force measured in mouse tracheal rings after incubation with vehicle or IL-13 (100 ng/mL) for 12 hours, followed by UNC3230 (200 nM) for 1 hour, with a range of concentrations of methacholine. n=5-6 rings per group. **e,** PIP5K1γ activity normalized to vehicle in HASM after treatment with vehicle or IL-13 (100 ng/mL) in presence of vehicle or Y-27632 (100 μM) for 12 hours, followed by lysis and IP with anti-PIP5K1γ antibody. n=6 biological replicates per group. **f,** Representative histograms for activated β1 integrin in HASM cells treated with vehicle, Mn^2+^ (1 mM) for 20 minutes, IL-13 (100 ng/mL) or IL-17A (100 ng/mL) in the presence of vehicle or Y-27632 (100 μM) for 12 hours, followed by labeling with HUTS-4 antibody and secondary conjugated to allophycocyanin (APC). **g,** Representative immunoprecipitation from lysates of mouse trachea after pulldown with rabbit IgG or PIP5K1γ antibody followed by immunoblot (IB) for Rho kinase. IB of PIP5K1γ was performed to confirm enrichment. Results representative of 3 biological replicates for **a**, **f**, and **g**. Data are mean ± s.e.m. for **b-e**. 2-way ANOVA with repeated measures, Tukey’s multiple-comparison test for **b** and **d**. 2-tailed t-test for **c**. 2-way ANOVA with Tukey’s multiple-comparison test for **e**. *P<0.05, **P<0.01, ***P<0.001, ****P<0.0001. #P<0.05, ####P<0.0001 vs IL-13 for **d**.

Importantly, upon activation of muscarinic receptors by methacholine, phospholipase C is activated to hydrolyze PIP_2_ to generate inositol 1,4,5-triphosphate (IP_3_), which directly mediates cytosolic Ca^2+^ release from the sarcoplasmic reticulum leading to muscle contraction (22). To determine whether our observed effects on force transmission were independent of IP_3_-related effects on contraction, we used potassium chloride (KCl) to directly activate voltage-gated calcium channels to allow for influx of extracellular calcium to initiate muscle contraction. diC16-PIP_2_ enhanced force transmission in mouse tracheal rings in response to increasing doses of KCl compared to rings treated with the carrier vehicle, suggesting that PIP_2_ can enhance force transmission in a manner independent of IP_3_ (**Supplemental Fig. 3a**).

The synthesis of PIP_2_ is primarily regulated by type I phosphatidylinositol 4-phosphate 5-kinases (PIP5K1) (23). In mammalian tissue there are three PIP5K1 isoforms (α, β, γ). We focused on PIP5K1γ because it is the major lipid kinase in airway smooth muscle (24, 25), has a focal adhesion targeting domain, and is recruited by talin which in turn binds to the integrin β subunit resulting in integrin activation (26, 27). To test the hypothesis that cytokines modulate PIP5K1γ activity, we measured PIP5K1γ activity after immunoprecipitating the enzyme from lysates of HASM cells treated with vehicle or IL-13. After normalization for protein concentration, PIP5K1γ kinase activity was higher in samples treated with IL-13 compared to vehicle (**Fig. 4c**). The enhanced PIP5K1γ activity induced by IL-13 could be inhibited by UNC3230, a small molecule inhibitor with good specificity for PIP5K1γ (**Supplemental Fig. 3b**) (28). Treatment of HASM cells with UNC3230 also inhibited integrin activation induced by IL-13 (**Supplemental Fig. 3c**) but had no effect when integrin activation was short-circuited by external addition of Mn^2+^ prior to flow cytometry (**Supplemental Fig. 5a**). Furthermore, while treatment with UNC3230 had no effect on force transmission in mouse tracheal rings at baseline compared to vehicle, UNC3230 completely abrogated the enhanced force transmission in rings treated with IL-13 (**Fig. 4d**). Taken together, these data support the hypothesis that the enhanced force transmission induced by pro-inflammatory cytokines is at least in part dependent on local modulation of PIP5K1 γ activity and PIP_2_ production.

PIP5K1-mediated synthesis of PIP_2_ is regulated by RhoA and its effector Rho kinase (29–31). To determine whether Rho kinase could regulate cytokine-mediated activation of integrins, we exposed HASM cells to IL-13 or IL-17A in the presence of vehicle or Y-27632, a small molecular inhibitor of Rho kinase. Y-27632 abrogated the enhanced PIP5K1γ activity induced by IL-13 in HASM cells (**Fig. 4e**). Furthermore, Y-27632 eliminated the integrin activation induced by IL-13 or IL-17A (**Fig. 4f**) but had no effect when integrins were externally activated by Mn^2+^ in flow cytometry (**Supplemental Fig. 5b**). We also found that Rho kinase precipitated with PIP5K1γ pulldown in lysates from mouse trachea (**Fig. 4g**). These findings reinforce the idea that pro-inflammatory cytokines activate β1 integrins through modulation of PIP5K1γ activity in a manner dependent on RhoA/Rho kinase.

### IL-13 and IL-17A modulate integrin activation through NF-kB

Expression of RhoA and its effector Rho kinase are known to be upregulated by a variety of pro-inflammatory cytokines including IL-4, IL-13, IL-17A, and TNF-α (32–34). We confirmed that RhoA and Rho kinase levels are increased after mouse tracheal strips are exposed to IL-13 or IL-17A compared to vehicle (**Supplemental Fig. 4a-b**). To confirm whether synthesis of new proteins was an important factor in the regulation of cytokine-mediated integrin activation, we first determined that HASM cells treated with IL-13 or IL-17A for 1 hour did not have an increase in active β1 integrin (**Fig. 5a**). Furthermore, treatment with the protein synthesis inhibitor cycloheximide eliminated the increase in active β1 integrin observed after 12 hours of treatment with either IL-13 or IL-17A (**Fig. 5b**).

**Figure 5:**
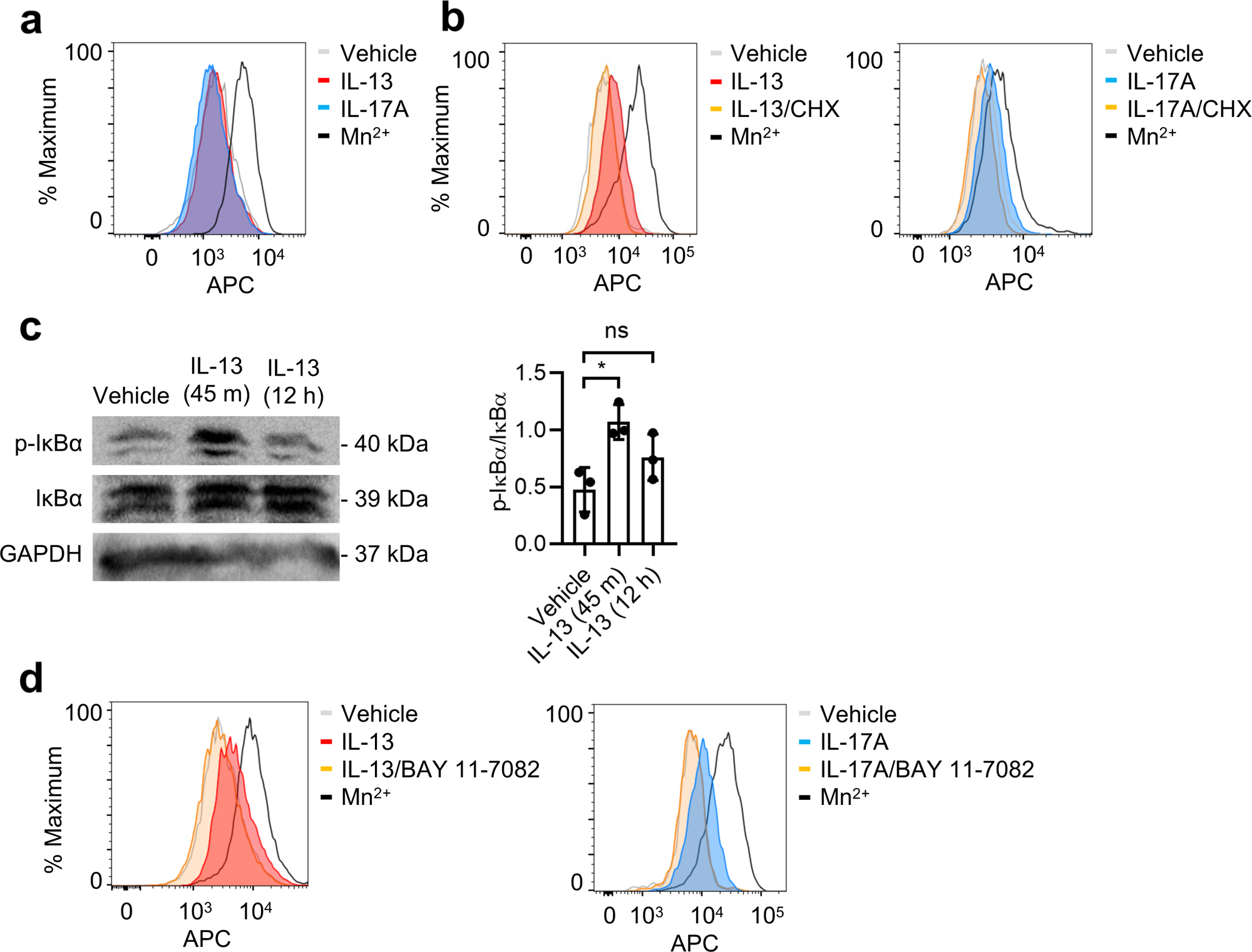
IL-13 and IL-17A modulate integrin activation through NF-κB. **a-b,** Representative histograms for active β1 integrin in HASM cells treated with vehicle, IL-13 (100 ng/mL), IL-17A (100 ng/mL) for (**a**) 1 hour or (**b**) 12 hours in the presence of cycloheximide (CHX, 15 ug/mL), or Mn^2+^ (1 mM) for 20 minutes followed by labeling with antibody specific for activated β1 integrin (HUTS-4) and secondary conjugated to allophycocyanin (APC). **c,** Representative western blot and densitometry for HASM treated with vehicle or IL-13 (100 ng/mL) for 45 minutes or 12 hours followed by histamine (100 μM) for 15 minutes followed by lysis, separation by SDS-PAGE, and transfer to membrane probed with antibodies to phospho- and total-IκBα and GAPDH. n=3 samples per condition. **d,** Representative histograms for active β1 integrin in HASM cells treated with vehicle, Mn^2+^ (1 mM) for 20 minutes, IL-13 (100 ng/mL) or IL-17A (100 ng/mL) in the presence of vehicle or BAY 11-7082 (2 μM) for 12 hours, followed by labeling with antibody specific for activated β1 integrin (HUTS-4) and secondary conjugated to allophycocyanin (APC). Results representative of 3 biological replicates for **a**, **b**, and **d**. Data are mean ± s.e.m. for **c**. 2-way ANOVA with Tukey’s multiple-comparison test for **c**. *P<0.05, ns=not significant.

To trigger transcriptional activation of new proteins, IL-13 binds to its receptor IL-13Rα1, which complexes with the IL-4Rα to induce phosphorylation of STAT6. STAT6 has also been shown to interact with NF-kB p50 and p65 to enhance its DNA binding affinity and transactivating ability (35). IL-17A triggers transcriptional activation through Act1-induced ubiquitylation of TRAF6 to activate MAPK, C/EBPβ, and NF-kB pathways. Both STAT6 and NF-kB are downstream targets of other pro-inflammatory cytokines in the asthmatic airway including IL-4 and TNF-α, and both have been shown to bind to the promoter regions of the RhoA gene (34).

IL-13 has also been shown to activate NF-kB in smooth muscle (36), and tissue expression of IL-13 can induce local NF-kB activity in vivo (37). We found that treatment of HASM cells with IL-13 induces NF-kB signaling as shown by a transient increase in phosphorylation of the IkBα complex (**Fig. 5c**). We therefore hypothesized that NF-kB may play a common downstream role in regulating integrin activation induced by multiple cytokines. Indeed, treatment with the NF-kB inhibitor BAY 11-7082, which inhibits the phosphorylation of IkBα, eliminated the integrin activation induced by IL-13 or IL-17A in HASM cells (**Fig. 5d**), but had no effect when integrins were externally activated by Mn^2+^ (**Supplemental Fig. 5c**) as assessed by flow cytometry. In contrast, inhibition of another downstream target of IL-17A, the MAPK pathway using the inhibitor SB 203580, did not affect integrin activation induced by either IL-13 or IL-17A (**Supplemental Fig. 6a-b**). Taken together, these results suggest that IL-13 and IL-17A activate integrins through NF-kB mediated upregulation of RhoA and Rho kinase.

## DISCUSSION

Integrin activation is central to cell-matrix dependent processes in both normal and pathologic conditions. In circulating cells such as leukocytes and platelets, integrins are predominantly in an inactive confirmation but are activated within seconds when exposed to chemokines that bind heterotrimeric G-protein coupled receptors (GPCR) at the cell surface (38–40). Rapid activation of integrins in circulating cells is essential to allow leukocyte extravasation to sites of inflammation (41) and platelet adherence to the vessel wall during injury (42). In stationary cells, integrin activation has generally been assumed to be less relevant, because the extended-open conformation is thought to predominate. Our findings add substantial depth to this supposition by demonstrating that while adherent muscle does indeed have substantial active β1 integrin, this can be further augmented by stimulation of receptors in the type I cytokine and IL-17 families, by IL-13 and IL-17A, in a manner dependent on NF-kB resulting in increased expression of RhoA and its effector Rho kinase. Induction of Rho kinase enhances PIP5K1γ activity resulting in local generation of PIP_2_ and conformational changes in β1 integrins. More importantly, these findings suggest that integrin activation itself, even in the absence of cytokine, has disease-relevant effects in chronic airways diseases such as asthma by enhancing force transmission in contracting muscle. Based on these observations, we propose a cascade of molecular events that regulate activation of β1 integrins by type I and IL-17 cytokine receptor stimulation (**Fig. 6**).

**Figure 6:**
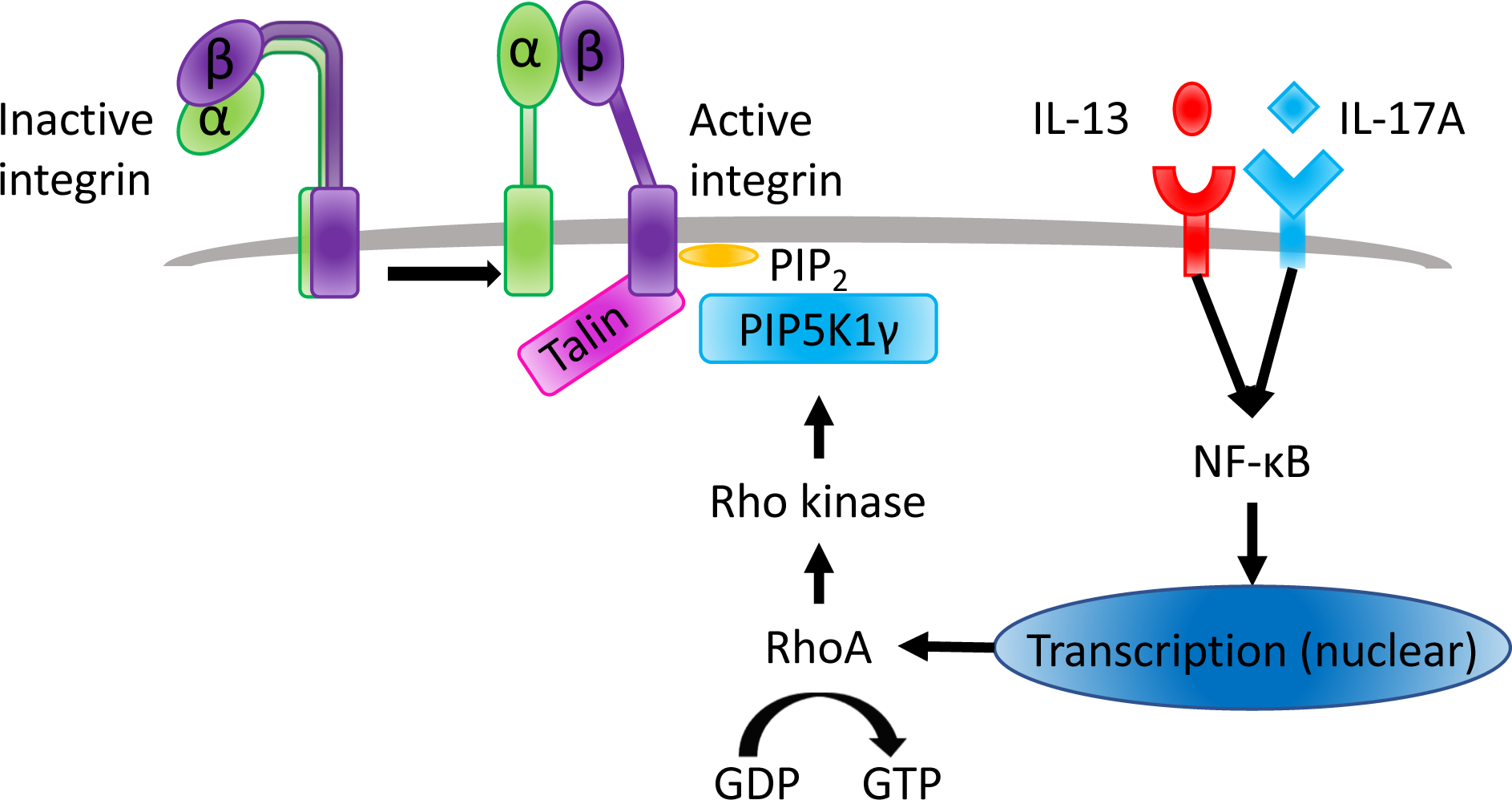
**Cytokines activate β1 integrins in smooth muscle by upregulating RhoA/Rho kinase to enhance PIP5K1γ kinase activity.**

Molecular pathways leading to integrin activation have been best studied in circulating cells, and generally involve three steps: (1) chemokine-mediated GPCR activation, (2) induction of Ras homolog family (Rho) GTPases, and (3) recruitment of talin to induce conformational change in the integrin. In addition to chemokine-induced GPCR activation, integrin affinity can also be modulated by stimulation of receptor tyrosine kinases and Toll-like/IL1 receptors (43, 44). Our results extend these observations further by demonstrating that entirely different classes of cytokine receptors, type I cytokine receptors activated by IL-13 and IL-17 receptors activated by IL-17A, are also capable of inducing integrin activation in adherent muscle, underscoring the structural diversity of cytokine receptors that can modulate integrin responses in tissue. While stimulation by different receptors may lead to activation of pathways that ultimately converge on integrin effector molecules, our data suggest that the timescale required for integrin activation can vary dramatically, with type I cytokine and IL-17 receptor stimulation requiring hours in contrast to the sub-second timescale required for chemokine-induced GPCR activation. As chemokine- and toll-like/IL1 receptor stimulation can also trigger multivalent clustering of integrins (40, 44), further investigation into whether IL-13 and IL-17A can induce changes in lateral mobility of integrins is necessary for a comprehensive understanding of how these cytokines influence overall integrin avidity. Nevertheless, our results have identified a distinct role for type I cytokine and IL-17 receptors in regulating integrin affinity in adherent cells.

From a mechanistic standpoint, classical chemokine-induced GPCR activation results in G protein-mediated activation of Rho GTPases. Gα activates Ras homolog family member A (RhoA)-mediated signaling modules (29, 45, 46) while Gβγ activates Ras-related C3 botulinum toxin substrate 1 (Rac1) and phospholipase C with associated intracellular Ca^2+^ flux (47) leading to activation of Ras-related protein 1 (Rap1) signaling modules (48, 49). Notably, activation of downstream signaling modules appears to be stimulus-, integrin-, and cell type-specific. For example, in lymphocytes CCL25 and CXCL10 stimulation result in distinct binding affinities of integrin α4β7 for MAdCAM-1 and VCAM-1 through differential activation of signaling modules (50). Furthermore, in neutrophils, macrophages, and conventional T cells Rap1-mediated activation of β2 integrins depends on the effector Rap1-GTP-interacting-adaptor molecule (RIAM), whereas RIAM appears to be dispensable for β1 and β3 integrin activation in leukocytes and platelets (51–54). Thus, while the distal molecular event of talin-induced integrin activation is likely conserved across cell types, it is reasonable to speculate that cell-specific differences in expression of signaling molecules or activation of signaling pathways allow for tissue-specific tuning of integrin activation in response to inflammatory stimuli. In this context, we identify a disease-relevant example of this activation heterogeneity in adherent smooth muscle with potential therapeutic relevance to chronic airways disease. Our findings linking cytokine-induced β1 integrin activation in HASM with modulation of RhoA and PIP5K1γ activity are concordant with prior studies in lymphocytes demonstrating that RhoA and PIP5K1γ can control specific modalities of chemokine-induced β2 integrin activation (29, 45, 55), and underscore the central role of Rho GTPases in regulating integrin activation. The dependence of cytokine-induced activation of β1 integrins on NF-kB and Rho kinase are noteworthy additions that may be specific to the specific cytokine receptor and cell type.

Another significant finding of our study is that the kinase activity of PIP5K1γ is enhanced by RhoA/Rho kinase, resulting in the recruitment, activation, and interaction of talin with the cytoplasmic tail of the integrin β-subunit. Noting the major source of PIP_2_ in airway smooth muscle is from PIP5K1γ (24), it is reasonable to hypothesize that our observed increases in β1 integrin activation are due to local increases in PIP_2_ which recruit and activate talin to induce conformational changes in the integrin (19, 20). Indeed, our finding that PIP_2_ enhances force transmission induced by voltage-gated ion channel stimulation (and independent of IP_3_-induced mobilization of calcium stores) also emphasizes the importance of PIP_2_ in recruiting talin and activating integrins. Recent studies have suggested that the 90-kD splice variant of PIP5K1γ (PIP5K1γ-90) can directly interact with the FERM (four-point-one-protein/ezrin/radixin/moesin) domain of talin (26). It therefore remains a possibility that PIP5K1γ may induce conformational changes in β1 integrins through direct interaction with talin.

Mechanical forces also play an important role in stabilization of nascent integrin adhesions (56–58). For example, tensile force has been shown to recruit paxillin and stretch talin to enhance vinculin binding, thus enhancing focal adhesion progression (59, 60). In airway smooth muscle, stabilization of adhesomes by tensile force plays a vital role in formation of the rigid network used to transmit force generated by sliding actin filaments across integrin-extracellular matrix tethers ultimately resulting in deformation of the airway lumen. In the setting of chronic inflammatory disorders of the airways, cytokine-mediated activation of surface β1 integrins is likely further stabilized by the exaggerated tensile force generated by hypercontractile smooth muscle. Our immunostaining of asthmatic airways demonstrating enhanced active β1 integrin in the smooth muscle reinforces this idea and spotlights a thus far unappreciated contributor to the exaggerated force transmission in asthmatic airways. It should also be noted that the effects of specific β1 heterodimer activation on force transmission may be varied. Our prior work with integrins α5β1 and α2β1 suggest that engagement of these integrins enhances force transmission and resultant airway narrowing (7, 8). On the other hand, integrin α9β1 appears to serve as a brake on airway contraction by reducing the activity of PIP5K1γ through its association with the polyamine catabolizing enzyme SSAT (25). The data we present here do not address specific integrin heterodimer effects but support the idea that on balance β1 integrin activation induced by IL-13 and IL-17A enhances force transmission in contracting muscle.

In summary, we found that stimulation of type I cytokine and IL-17 receptors by IL-13 and IL-17A, respectively, induces physiological activation of β1 integrins in adherent cells by enhancing PIP5K1γ activity in a manner dependent on NF-kB, RhoA, and its effector Rho kinase. This process requires hours, in marked distinction to the sub-second timescale required for chemokine-induced GPCR activation in circulating cells. Nevertheless, our findings provide crucial insight into the importance of integrin activation in adherent cells, suggest that the degree of integrin activation has functional effects on enhancing force transmission, and raise the possibility that this pathway could be therapeutically targeted to alleviate airway narrowing in inflammatory disorders of the airways.

## METHODS

### Reagents

GAPDH, phospho-IkBα, total IkBα, and Rho kinase antibodies were purchased from Cell Signaling Technology. PIP5K1γ antibody was purchased from Abcam. α-smooth muscle actin antibody was purchased from Sigma. Integrin antibodies used were monoclonal anti-active integrin β1 (HUTS-4, Millipore), activating integrin β1 (TS2/16, Invitrogen). c15 was purchased from Tocris. β1-CHAMP was synthesized in our laboratory (16). Manganese was purchased from Fisher. Y-27632 was purchased from Sigma. SB 203580 and BAY 11-7082 were purchased from MedChemExpress. UNC3230 was purchased from Cayman Chemical. PI(4,5)P2 diC16 was purchased from Echelon Biosciences and used with carrier 1 per manufacturer instructions. Cycloheximide was purchased from Research Products International (RPI).

### Cells

HASM cells and media were purchased from Lonza and cultured according to the vendor’s instructions. Cells were used between passage 5 and 10.

### Mice

Mice used for all experiments were in a C57BL/6 background, 6-10 weeks old, and housed under specific pathogen-free conditions in the Animal Barrier Facility at UCSF.

### Cell Adhesion assay

96-well flat-bottomed tissue culture plates (Earthox) were coated with varying concentrations of rat tail collagen I (Sigma) or fibronectin (Sigma) for 1 h at 37° C. After incubation, wells were washed with PBS, then blocked with 1% BSA at 37° C for 1 h. Control wells were filled with 1% BSA. HASM cells were detached using 10 mM EDTA and re-suspended in serum-free DMEM before plating 50,000 cells per well. The plates were centrifuged at 300 rpm for 5 min before incubation for 1 h at 37° C in humidified 5% CO_2_. Non-adherent cells were removed by centrifugation (top side down) at 300 rpm for 5 min. Attached cells were stained with 0.5% crystal violet and the wells were washed with PBS. The relative number of cells in each well was evaluated after solubilization in 40 µL of 2% Triton X-100 by measuring absorbance at 595 nm in a microplate reader (Bio-Rad Laboratories). All determinations were carried out in triplicate.

### Flow cytometry

HASM cells were harvested with 0.25% trypsin-EDTA, washed twice with PBS and re-suspended in FACS buffer (HEPES buffered saline, supplemented with 10% serum and 1% BSA). 5 x 10^5^ cells were incubated with a primary antibody at 4° C for 30 min in the dark. Cells were then washed, re-suspended in FACS buffer and incubated with secondary goat anti-mouse antibody conjugated to allophycocyanin (APC; Jackson ImmunoResearch). Cells were then washed, re-suspended in 2.5% serum, and analyzed on a BD FACSCantoll. Antibodies were used at 10 µg/mL. Integrin activation was achieved with exposure to human IL-13 or IL-17A (100 ng/mL; Peprotech) for 12 h or Mn^2+^ (1 mM) for 20 minutes before incubating with primary antibody. All analyses were performed after gating for live cells.

### Collagen force transmission assay

Collagen gels were prepared from on ice with PBS, DMEM, 10 µM NaOH, and Type I collagen (Corning) to achieve collagen concentration 2 mg/mL. HASM cells were harvested, suspended in DMEM, and mixed in 1:1 volume ratio with collagen at a concentration of 2.5 x 10^5^ cells per ml. Gels were polymerized in 24 well plates at 37°C for 90 minutes followed by mechanical release and overnight incubation in DMEM. Images were taken at baseline and 20 minutes after the addition of 100 µM histamine. Changes in collagen gel area was analyzed using ImageJ and expressed as a percentage of baseline.

### Measurement of tracheal smooth muscle contractility

Tracheal ring contraction studies were performed as described previously (25, 61). Briefly, after incubation with the indicated treatment, rings were equilibrated under 0.5 gm of applied tension, contracted with 60 mM KCl, and only rings that generated more than 1 mN of force were analyzed. After re-equilibration, contractile responses were evaluated to increasing doses of methacholine or KCl (Sigma-Aldrich). For analysis of suppressive effects on cytokine-induced contractility, rings were treated with human IL-13 (100 ng/mL; Peprotech) or human IL-17A (100 ng/mL; Peprotech) for 12 h at 37°C with 5% CO_2_. For human bronchial rings, lung tissue was obtained from lung transplant donors. Bronchi, 5-8 mm in diameter, were dissected free of connective tissue and cut into 4-mm-thick rings. Rings were stored and assessed as above, except a resting tension of 1g was applied, and rings were first contracted with 120 mM KCl after equilibration for 2 hours, and only the rings that generated more than 2 mN of force were used for experiments.

### Immunostaining

Human bronchial rings were embedded in Optimal Cutting Temperature compound, and 5-µm sections were prepared using a Leica Cryostat (CM 1850). The sections were fixed with acetone for 5 minutes at -20° C, blocked with 10% serum, and incubated overnight with a mouse IgG2b monoclonal antibody (clone HUTS-4) directed against active β1 integrin and α-smooth muscle actin (clone 1A4, FITC-conjugated; Sigma), followed by 1 hour incubation with anti-mouse IgG2b Alexa-647. Sections were mounted in mounting medium with DAPI and visualized with a Leica DM5000B epifluorescent microscope. Endobronchial biopsy specimens were obtained from patients with asthma based on positive bronchial provocation testing as well as healthy non-smoking controls. Biopsy samples were embedded in Optimal Cutting Temperature compound and frozen in liquid nitrogen in the Airway Tissue Bank at UCSF. The tissue bank is approved by the UCSF Committee on Human Research. Samples were processed as above.

### Immunoblots

Smooth muscle dissected from mouse tracheas or HASM cells were homogenized in lysis buffer (50 mM Tris-HCl, pH 7.5, 10 mM MgCl_2_, 150 mM NaCl, 1% Triton X-100, 10 mM NaF, 1 mM Na_3_VO_4_) with protease and phosphatase inhibitor cocktail (Thermo). Lysates were centrifuged and supernatant resolved by SDS-PAGE. For immunoprecipitation, lysed samples were incubated with PIP5K1γ antibody and protein G sepharose beads for 4 hours at 4°C with rotation (70 rpm). Samples were washed 4 times with 1 mL lysis buffer and eluted with reducing sample buffer then resolved by SDS-PAGE. After transfer to a polyvinylidene difluoride membrane (Millipore), membranes were blocked for 1 h with 5% BSA in Tris buffered saline with Tween-20, incubated at room temperature for 2 h with primary antibodies, washed in Tris buffered saline with Tween-20, incubated for 1 h with peroxidase-conjugated secondary antibody, washed in Tris buffered saline with Tween-20, and developed with ECL plus (Perkin Elmer) prior to chemiluminescence detection (Bio-Rad Laboratories). All quantitative densitometry was calculated with ImageJ.

### PIP5K1Y kinase activity assay

HASM cells were lysed in IP buffer (50 mM Tris-HCl, pH 7.5; 150 mM NaCl; 0.01% Triton X-100; 5.0 mM NaF; 2 mM Na_3_VO_4_; 1 mM EDTA; 0.1 mM EGTA; 10% glycerol; and proteinase inhibitor cocktail), centrifuged, and incubated with Protein A-sepharose and rabbit anti-PIP5K1γ antibody (ab109192, Abcam) at 4°C overnight. Sepharose beads were then washed 3 times with IP buffer, with final wash containing 0.01% Triton X-100. PIP5K1γ kinase activity was assayed following a previously described method for lipid kinases (62). Briefly, kinase activity was assayed at room temperature in 20 µl buffer substrate solution and 20 µl IP protein. Reaction conditions consisted of 30 mM Tris (pH 7.4), 1.2 mM MgCl_2_, 0.3 mg/ml BSA, 60 µM dithiothreitol, 0.06 mg/ml PI4P/DOPS (1:1; Avanti Polar Lipids), 3.5 µM ATP, and 22 µCi [γ-^32^P] ATP (Perkin Elmer). The kinase reaction was started by the addition of ATP and was terminated after 3 hours by spotting 4 µl of the reaction onto 0.2 µm nitrocellulose membrane. When spotting was complete, the membrane was dried under a heat lamp for 5 minutes, washed once with 200 ml of 1% phosphoric acid in 1 M NaCl for 30 seconds, followed by 6 washes for 5 minutes. The membrane was dried for 20 minutes, exposed for 16 hours on a phosphor screen, and scanned using the Typhoon (GE Healthcare). Spot intensity was quantified by Image Studio Lite. PIP5K1γ activity was normalized by protein concentrations in lysates and expressed as a percentage of control.

### Statistics

The statistical significance of the differences within or between multiple groups was calculated with a 1-way or 2-way ANOVA, with repeated measures of variance for related samples, and when differences were statistically significant (P≤0.05), this was followed with a Sidak or Tukey *t* test for subsequent pairwise analysis. All calculations were performed using Prism (GraphPad Software).

### Study Approval

All mice were housed in a specific pathogen-free animal facility at the University of California, San Francisco. Animals were treated according to protocols that were approved by the Institutional Animal Care and Use Committee at UCSF in accordance with the US National Institutes of Health (NIH) guidelines.

## AUTHOR CONTRIBUTIONS

A.S. designed the research studies. U.N. and A.S. performed most of the in vitro and ex vivo experiments. Y.S. performed and analyzed kinase activity assays. H.J. performed chemical synthesis. U.N. and A.S. collected and analyzed the data. P.W. provided human samples and technical support for immunostaining. K.S., W.D., D.S. provided reagents, equipment, and conceptual advice. U.N. and A.S. wrote the manuscript with input from all authors.

## ACKNOWLEDGEMENTS

The authors thank all members of the Sandler Asthma Basic Research Group for helpful discussions and critical review. This work was supported by NIH grants K08 HL124049 and R61 HL163725 to A. Sundaram, P01 HL146373 and R35 GM122603 to H. Jo.

## Notes

### Competing Interest Statement

The authors have declared no competing interest.

## REFERENCES

1. Fehrenbach H, Wagner C, Wegmann M. Airway remodeling in asthma: what really matters. Cell Tissue Res. 2017;367(3):551–569.

2. Wills-Karp M, et al. Interleukin-13: central mediator of allergic asthma. Science. 1998;282(5397):2258–2261.

3. Munitz A, et al. Distinct roles for IL-13 and IL-4 via IL-13 receptor alpha1 and the type II IL-4 receptor in asthma pathogenesis. Proc Natl Acad Sci U S A. 2008;105(20):7240–7245.

4. Bullens DM, et al. IL-17 mRNA in sputum of asthmatic patients: linking T cell driven inflammation and granulocytic influx? Respir Res. 2006;7(1):135.

5. Al-Ramli W, et al. TH17-associated cytokines (IL-17A and IL-17F) in severe asthma. Journal of Allergy and Clinical Immunology. 2009;123(5):1185–1187.

6. Hynes RO. Integrins: bidirectional, allosteric signaling machines. Cell. 2002;110(6):673–687.

7. Sundaram A, et al. Targeting integrin α5β1 ameliorates severe airway hyperresponsiveness in experimental asthma. J Clin Invest. 2017;127(1):365–374.

8. Liu S, et al. Integrin α2β1 regulates collagen I tethering to modulate hyperresponsiveness in reactive airway disease models. J Clin Invest. [published online ahead of print: May 6, 2021]. 10.1172/JCI138140.

9. Schumacher S, et al. Structural insights into integrin α5β1 opening by fibronectin ligand. Science Advances. 2021;7(19):eabe9716.

10. Xiong JP, et al. Crystal structure of the extracellular segment of integrin alpha Vbeta3. Science. 2001;294(5541):339–345.

11. Takagi J, et al. Global conformational rearrangements in integrin extracellular domains in outside-in and inside-out signaling. Cell. 2002;110(5):599–511.

12. Luque A, et al. Activated Conformations of Very Late Activation Integrins Detected by a Group of Antibodies (HUTS) Specific for a Novel Regulatory Region(355-425) of the Common β1 Chain (∗). Journal of Biological Chemistry. 1996;271(19):11067–11075.

13. Chen J, Salas A, Springer TA. Bistable regulation of integrin adhesiveness by a bipolar metal ion cluster. Nat Struct Mol Biol. 2003;10(12):995–1001.

14. Kim M, Carman CV, Springer TA. Bidirectional Transmembrane Signaling by Cytoplasmic Domain Separation in Integrins. Science. 2003;301(5640):1720–1725.

15. Fong KP, et al. Directly Activating the Integrin αIIbβ3 Initiates Outside-In Signaling by Causing αIIbβ3 Clustering. J Biol Chem. 2016;291(22):11706–11716.

16. Mravic M, et al. De novo designed transmembrane peptides activating the α5β1 integrin. Protein Eng Des Sel. 2018;31(5):181–190.

17. Fong KP, et al. Visualization of Platelet Integrins via Two-Photon Microscopy Using Anti-transmembrane Domain Peptides Containing a Blue Fluorescent Amino Acid. Biochemistry. 2021;60(21):1722–1730.

18. Song X, et al. A novel membrane-dependent on/off switch mechanism of talin FERM domain at sites of cell adhesion. Cell Res. 2012;22(11):1533–1545.

19. Goksoy E, et al. Structural basis for the autoinhibition of talin in regulating integrin activation. Mol Cell. 2008;31(1):124–133.

20. Martel V, et al. Conformation, localization, and integrin binding of talin depend on its interaction with phosphoinositides. J Biol Chem. 2001;276(24):21217–21227.

21. Dedden D, et al. The Architecture of Talin1 Reveals an Autoinhibition Mechanism. Cell. 2019;179(1):120–131.e13.

22. Hall IP. Second messengers, ion channels and pharmacology of airway smooth muscle. Eur Respir J. 2000;15(6):1120.

23. Stephens LR, Hughes KT, Irvine RF. Pathway of phosphatidylinositol(3,4,5)-trisphosphate synthesis in activated neutrophils. Nature. 1991;351(6321):33–39.

24. Chen H, et al. Effects of polyamines and calcium and sodium ions on smooth muscle cytoskeleton-associated phosphatidylinositol (4)-phosphate 5-kinase. Journal of Cellular Physiology. 1998;177(1):161–173.

25. Chen C, et al. Integrin α9β1 in airway smooth muscle suppresses exaggerated airway narrowing. J Clin Invest. 2012;122(8):2916–2927.

26. Di Paolo G, et al. Recruitment and regulation of phosphatidylinositol phosphate kinase type 1 gamma by the FERM domain of talin. Nature. 2002;420(6911):85–89.

27. Ling K, et al. Type Iγ phosphatidylinositol phosphate kinase targets and regulates focal adhesions. Nature. 2002;420(6911):89–93.

28. Wright BD, et al. The lipid kinase PIP5K1C regulates pain signaling and sensitization. Neuron. 2014;82(4):836–847.

29. Bolomini-Vittori M, et al. Regulation of conformer-specific activation of the integrin LFA-1 by a chemokine-triggered Rho signaling module. Nat Immunol. 2009;10(2):185–194.

30. Weernink PAO, et al. Activation of Type I Phosphatidylinositol 4-Phosphate 5-Kinase Isoforms by the Rho GTPases, RhoA, Rac1, and Cdc42*. Journal of Biological Chemistry. 2004;279(9):7840–7849.

31. Chong LD, et al. The small GTP-binding protein Rho regulates a phosphatidylinositol 4-phosphate 5-kinase in mammalian cells. Cell. 1994;79(3):507–513.

32. Kudo M, et al. IL-17A produced by αβ T cells drives airway hyper-responsiveness in mice and enhances mouse and human airway smooth muscle contraction. Nat Med. 2012;18(4):547– 554.

33. Chiba Y, et al. Interleukin-13 augments bronchial smooth muscle contractility with an up-regulation of RhoA protein. Am J Respir Cell Mol Biol. 2009;40(2):159–167.

34. Goto K, et al. The proximal STAT6 and NF-kB sites are responsible for IL-13- and TNF-α-induced RhoA transcriptions in human bronchial smooth muscle cells. Pharmacol Res. 2010;61(5):466–472.

35. Shen C-H, Stavnezer J. Interaction of Stat6 and NF-kB: Direct Association and Synergistic Activation of Interleukin-4-Induced Transcription. Mol Cell Biol. 1998;18(6):3395–3404.

36. Goto K, Chiba Y, Misawa M. IL-13 induces translocation of NF-kappaB in cultured human bronchial smooth muscle cells. Cytokine. 2009;46(1):96–99.

37. Chapoval SP, et al. Inhibition of NF-kappaB activation reduces the tissue effects of transgenic IL-13. J Immunol. 2007;179(10):7030–7041.

38. Lagarrigue F, Kim C, Ginsberg MH. The Rap1-RIAM-talin axis of integrin activation and blood cell function. Blood. 2016;128(4):479–487.

39. Schmid MC, et al. PI3Kγ stimulates a high molecular weight form of myosin light chain kinase to promote myeloid cell adhesion and tumor inflammation. Nat Commun. 2022;13:1768.

40. Constantin G, et al. Chemokines Trigger Immediate β2 Integrin Affinity and Mobility Changes: Differential Regulation and Roles in Lymphocyte Arrest under Flow. Immunity. 2000;13(6):759–769.

41. Ley K, et al. Getting to the site of inflammation: the leukocyte adhesion cascade updated. Nat Rev Immunol. 2007;7(9):678–689.

42. Bennett JS, Vilaire G. Exposure of platelet fibrinogen receptors by ADP and epinephrine. J Clin Invest. 1979;64(5):1393–1401.

43. Schmid MC, et al. Receptor Tyrosine Kinases and TLR/IL1Rs Unexpectedly Activate Myeloid Cell PI3Kγ, A Single Convergent Point Promoting Tumor Inflammation and Progression. Cancer Cell. 2011;19(6):715–727.

44. Maynard SA, et al. IL-1β mediated nanoscale surface clustering of integrin α5β1 regulates the adhesion of mesenchymal stem cells. Sci Rep. 2021;11(1):6890.

45. Giagulli C, et al. RhoA and Ç PKC Control Distinct Modalities of LFA-1 Activation by Chemokines: Critical Role of LFA-1 Affinity Triggering in Lymphocyte In Vivo Homing. Immunity. 2004;20(1):25–35.

46. Soede RDM, et al. Stromal Cell-Derived Factor-1-Induced LFA-1 Activation During In Vivo Migration of T Cell Hybridoma Cells Requires Gq/11, RhoA, and Myosin, as well as Gi and Cdc421. The Journal of Immunology. 2001;166(7):4293–4301.

47. Block H, et al. Gnb isoforms control a signaling pathway comprising Rac1, Plcβ2, and Plcβ3 leading to LFA-1 activation and neutrophil arrest in vivo. Blood. 2016;127(3):314–324.

48. Katagiri K, et al. RAPL, a Rap1-binding molecule that mediates Rap1-induced adhesion through spatial regulation of LFA-1. Nat Immunol. 2003;4(8):741–748.

49. Shimonaka M, et al. Rap1 translates chemokine signals to integrin activation, cell polarization, and motility across vascular endothelium under flow. J Cell Biol. 2003;161(2):417– 427.

50. Sun H, et al. Distinct Chemokine Signaling Regulates Integrin Ligand Specificity to Dictate Tissue-Specific Lymphocyte Homing. Developmental Cell. 2014;30(1):61–70.

51. Lafuente EM, et al. RIAM, an Ena/VASP and Profilin Ligand, Interacts with Rap1-GTP and Mediates Rap1-Induced Adhesion. Developmental Cell. 2004;7(4):585–595.

52. Klapproth S, et al. Loss of the Rap1 effector RIAM results in leukocyte adhesion deficiency due to impaired β2 integrin function in mice. Blood. 2015;126(25):2704–2712.

53. Stritt S, et al. Rap1-GTP–interacting adaptor molecule (RIAM) is dispensable for platelet integrin activation and function in mice. Blood. 2015;125(2):219–222.

54. Sun H, et al. Distinct integrin activation pathways for effector and regulatory T cell trafficking and function. Journal of Experimental Medicine. 2020;218(2):e20201524.

55. Laudanna C, Campbell JJ, Butcher EC. Role of Rho in Chemoattractant-Activated Leukocyte Adhesion Through Integrins. Science. 1996;271(5251):981–983.

56. Nordenfelt P, Elliott HL, Springer TA. Coordinated integrin activation by actin-dependent force during T-cell migration. Nature Communications. 2016;7. 10.1038/ncomms13119.

57. Li J, Springer TA. Integrin extension enables ultrasensitive regulation by cytoskeletal force. Proc Natl Acad Sci U S A. 2017;114(18):4685–4690.

58. Stricker J, et al. Spatiotemporal Constraints on the Force-Dependent Growth of Focal Adhesions. Biophysical Journal. 2011;100(12):2883–2893.

59. Yao M, et al. Mechanical activation of vinculin binding to talin locks talin in an unfolded conformation. Sci Rep. 2014;4:4610.

60. Opazo Saez A, et al. Tension development during contractile stimulation of smooth muscle requires recruitment of paxillin and vinculin to the membrane. American Journal of Physiology-Cell Physiology. 2004;286(2):C433–C447.

61. Chen C, Huang X, Sheppard D. ADAM33 Is Not Essential for Growth and Development and Does Not Modulate Allergic Asthma in Mice. Mol Cell Biol. 2006;26(18):6950–6956.

62. Knight ZA, et al. A membrane capture assay for lipid kinase activity. Nat Protoc. 2007;2(10):2459–2466.

