## supplement for "IL-13 and IL-17A Activate β1 Integrin through an NF-kB/Rho kinase/PIP5K1γ pathway to Enhance Force Transmission in Airway Smooth Muscle"

**Supplemental Figure 1**

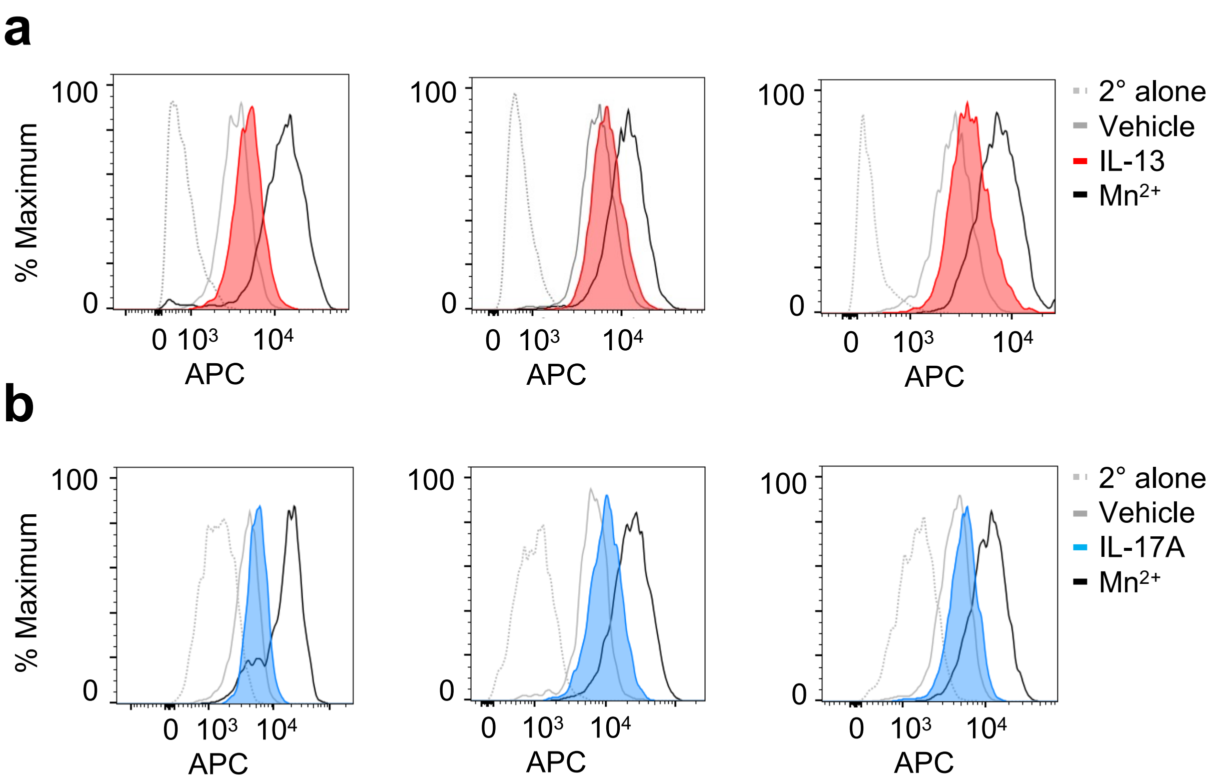


**Supplemental Figure 1: IL-13 and IL-17A enhance adhesion and activate β1 integrins.**
**a-b,** Histograms for activated β1 integrin in HASM cells treated with vehicle, $Mn^{2+}$ (1 mM) for 20 minutes, or (**a**) IL-13 (100 ng/mL) or (**b**) IL-17A (100 ng/mL) for 12 hours followed by labeling with antibody specific for activated β1 integrin (HUTS-4) and secondary (2°) conjugated to allophycocyanin (APC). Results are shown from 3 independent biological replicates for **a-b**.

**Supplemental Figure 2**

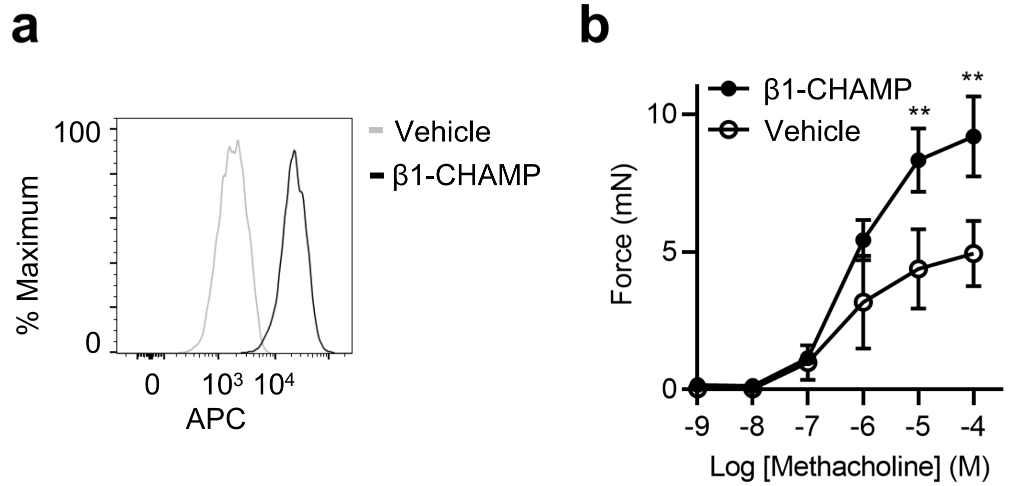


**Supplemental Figure 2: β1-CHAMP activates β1 integrins and enhances force transmission.**
**a,** Representative histogram for activated β1 integrin in HASM cells treated with vehicle or β1-CHAMP (10 μM) for 1 hour followed by labeling with antibody specific for activated β1 integrin (HUTS-4) and secondary conjugated to allophycocyanin (APC). Results representative of 3 biological replicates. **b,** Contractile force measured in mouse tracheal rings after incubation with vehicle or β1-CHAMP (10 μM) for 1 hour, with a range of concentrations of methacholine. n=3 rings per group. Data are mean ± s.e.m. for **b**. 2-way ANOVA with repeated measures, Tukey's multiple-comparison test for **b**. **P<0.01.

**Supplemental Figure 3**

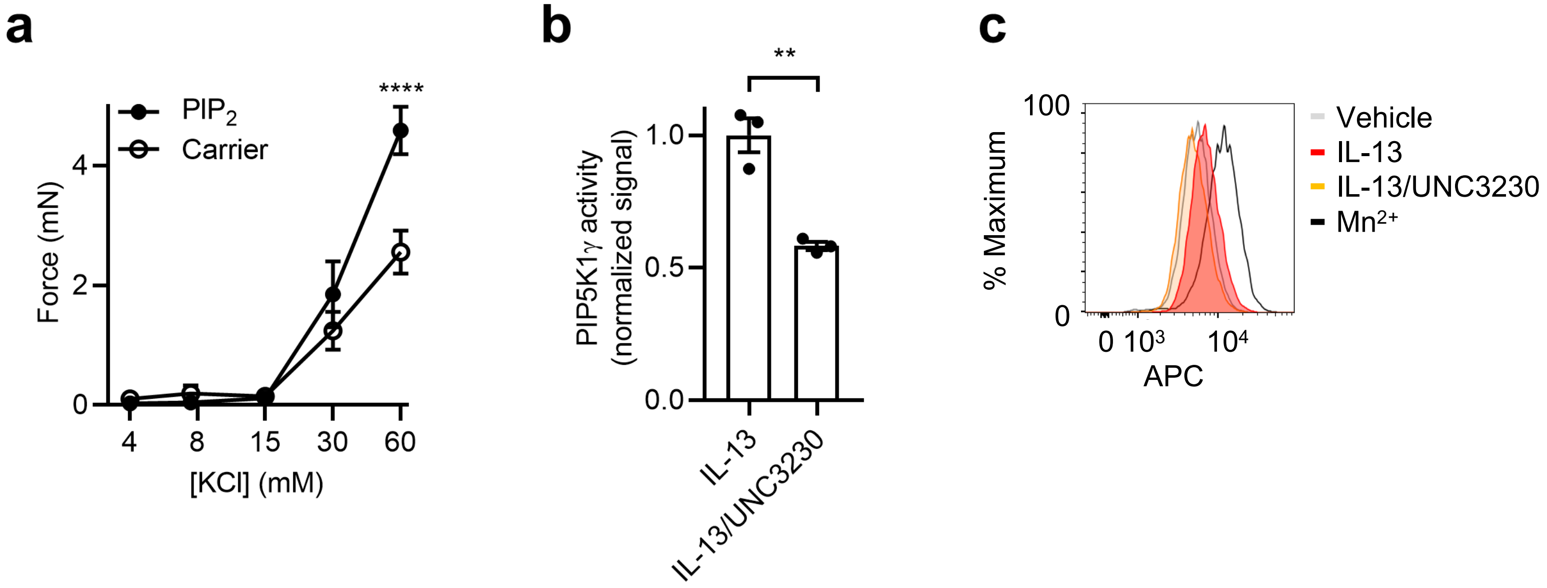


**Supplemental Figure 3: PIP5K1γ-mediated synthesis of $PIP_2$ activates β1 integrins and enhances force transmission.**

**a,** Contractile force measured in mouse tracheal rings after incubation with lipid carrier 1 in the presence or absence of diC16-$PIP_2$ (10 μM) for 1 hour, with a range of concentrations of KCl. n=12-13 rings per group. **b,** PIP5K1γ activity normalized to vehicle in HASM after treatment with IL-13 (100 ng/mL) in the presence of vehicle or Y-27632 (100 μM) for 12 hours, followed by lysis and IP with anti-PIP5K1γ antibody. n=3 biological replicates per group. **c,** Representative histogram for activated β1 integrin in HASM cells treated with vehicle, $Mn^{2+}$ (1 mM) for 20 minutes, or IL-13 (100 ng/mL) in the presence of vehicle or UNC3230 (200 nM) for 12 hours, followed by labeling with antibody specific for activated β1 integrin (HUTS-4) and secondary conjugated to allophycocyanin (APC). Results representative of 3 biological replicates. Data are mean ± s.e.m. for **a-b**. 2-way ANOVA with repeated measures, Tukey's multiple-comparison test for **a**. 2-tailed t-test for **b**. $^{**}P<0.01$, $^{****}P<0.0001$.

**Supplemental Figure 4**

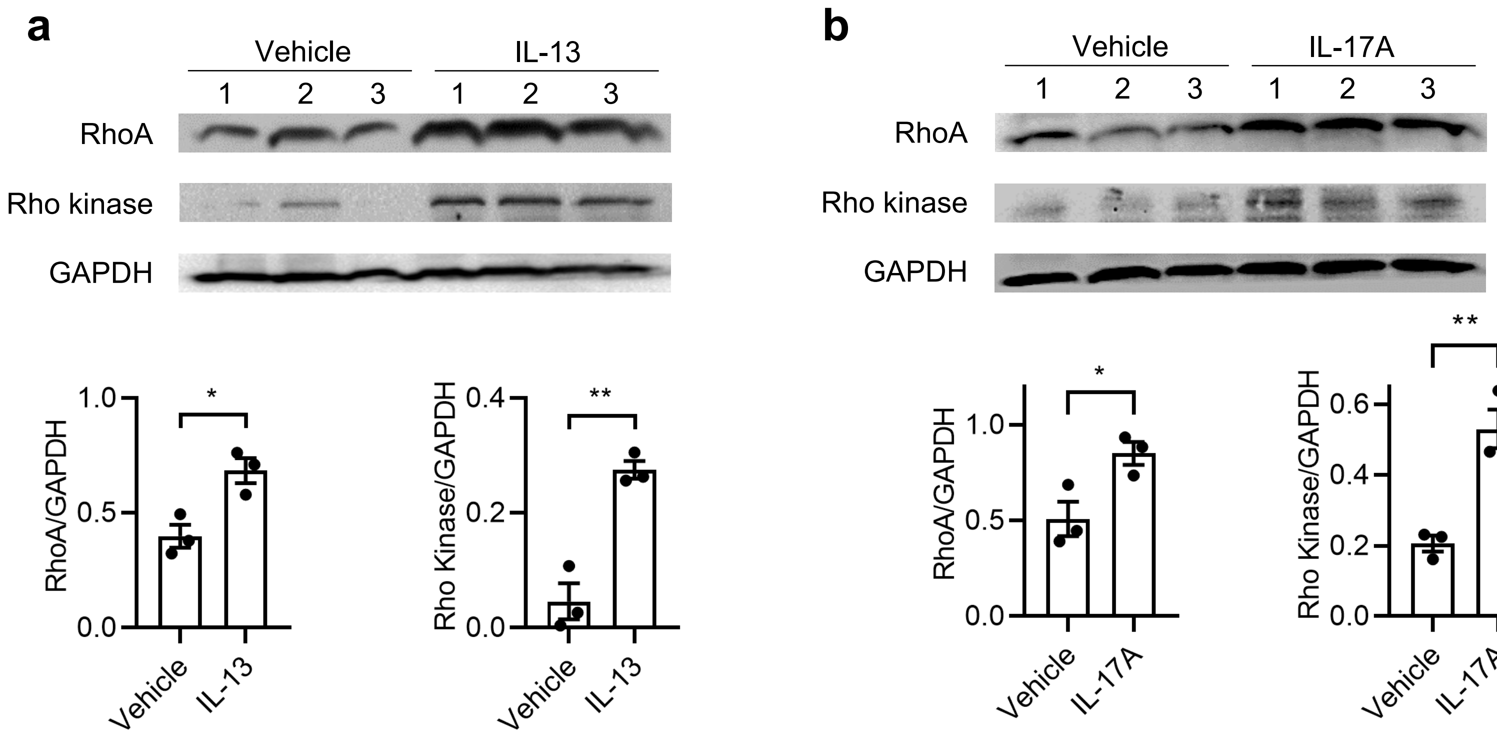


**Supplemental Figure 4: IL-13 and IL-17A activate RhoA and Rho kinase.**

**a-b,** Representative western blot and densitometry for mouse tracheal strips treated with vehicle, (**a**) IL-13 (100 ng/mL) or (**b**) IL-17A (100 ng/mL) for 12 hours followed by methacholine ($10^{-4}$ M) for 5 minutes followed by lysis, separation by SDS-PAGE, and transfer to membrane probed with antibodies to RhoA, Rho kinase, and GAPDH. n=3 tracheal strips per condition for **a-b**. Data are mean ± s.e.m. for **a-b**. 2-tailed t-test for **a-b**. *P<0.05, **P<0.01.

**Supplemental Figure 5**

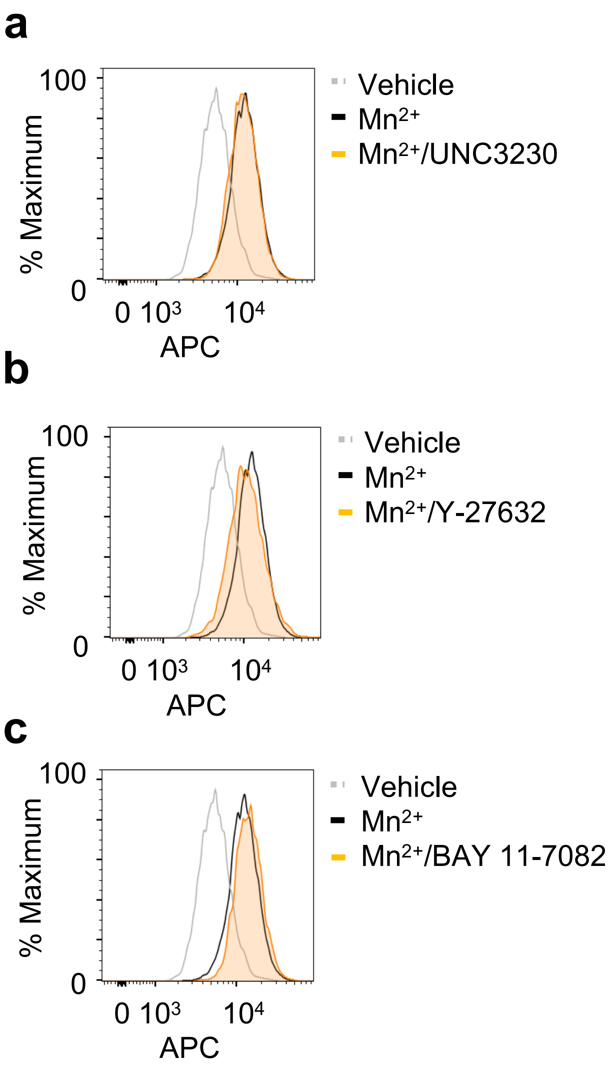


**Supplemental Figure 5: Inhibitors do not modulate external integrin activation.**
**a-c,** Representative histograms for activated β1 integrin in HASM cells treated with vehicle, (**a**) UNC3230 (100 μM), (**b**) Y-27632 (100 μM), or (**c**) BAY 11-7082 (2 μM) for 12 hours, then $Mn^{2+}$ (1 mM) for 20 minutes, followed by labeling with antibody specific for activated β1 integrin (HUTS-4) and secondary conjugated to allophycocyanin (APC). Results representative of 3 biological replicates for **a-c**.

### Supplemental Figure 6

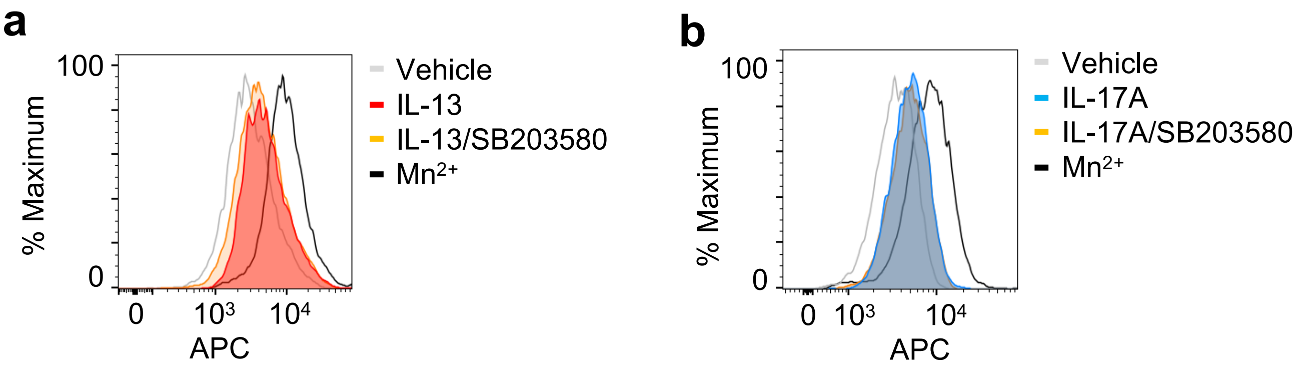


**Supplemental Figure 6: Cytokine-dependent integrin activation is independent of MAPK pathway.**
**a-b,** Representative histograms for activated β1 integrin in HASM cells treated with vehicle, $Mn^{2+}$ (1 mM) for 20 minutes, (**a**) IL-13 (100 ng/mL) or (**b**) IL-17A (100 ng/mL) in the presence of vehicle or SB203580 (100 μM) for 12 hours, followed by labeling with antibody specific for activated β1 integrin (HUTS-4) and secondary conjugated to allophycocyanin (APC). Results representative of 3 biological replicates for **a-b**.

**Supplemental Table 1**

| | Age (y) | Sex | $FEV_1$ L (%) | FVC L (%) | $FEV_1$/FVC |
|---|---|---|---|---|---|
| Healthy | 58 | Female | 2.44 (92) | 3.07 (114) | 0.79 |
| Healthy | 36 | Female | 3.11 (102) | 3.7 (117) | 0.84 |
| Healthy | 42 | Female | 4.27 (128) | 5.24 (104) | 0.81 |
| Asthma | 67 | Female | 1.81 (92) | 2.91 (92) | 0.62 |
| Asthma | 20 | Female | 3.36 (107) | 4.16 (103) | 0.81 |
| Asthma | 30 | Male | 3.78 (84) | 5.78 (128) | 0.65 |

**Supplemental Table 1: Demographic and Clinical Characteristics of Healthy and Asthmatic Patients.** Asthmatic patients were diagnosed based on positive bronchial provocation testing. $FEV_1$ = Forced Expiratory Volume in 1 second. FVC = Forced Vital Capacity.
